# Integrated model of the protein molecular clock across mammalian species

**DOI:** 10.1101/2025.01.14.633008

**Authors:** Anshul Saini, Dinara R. Usmanova, Dennis Vitkup

## Abstract

One of the foundational concepts in molecular evolution is the protein molecular clock [1-3], encapsulating the observation that proteins tend to accumulate amino acid substitutions at approximately constant and protein-specific rates over long evolutionary timescales. According to the neutral theory of molecular evolution, the majority of protein substitutions have neutral or nearly neutral effects on species fitness. The neutral theory suggests the existence of a strong generation time effect, positing that species with shorter generation times should accumulate protein substitutions faster per unit time compared to species with longer generation times. However, earlier studies failed to detect a significant generation time effect for amino acid substitutions in vertebrates, raising questions about core tenets of the neutral theory. In this study, we take advantage of the recently accumulated evolutionary and genomics data to investigate the scaling of substitution rates with generation time among many dozens of mammalian species and thousands of proteins. Our findings reveal a strong generation time effect both for synonymous and non-synonymous substitutions. We demonstrate that the empirically observed generation time effect is fully consistent with several mechanisms, such as functional and mutation rate selection. Building on these results, we develop an integrated model of protein evolution that incorporates changes in substitution rates due to variation in species generation times throughout their evolutionary history. Overall, our analysis shows that protein evolution is consistent with neutral or nearly neutral theories and explains time-like behavior of the protein molecular clock across deep phylogenetic lineages.

## Introduction

One of the major breakthroughs in understanding protein evolution has been the development of the neutral theory [4-6]. The neutral theory of molecular evolution suggests that the majority of amino acid substitutions in proteins are neutral or nearly neutral with respect to their effects on species’ fitness. The neutral theory forms a convenient null model of evolution at the molecular level [7-9] and provides mechanistic explanations to multiple empirical observations. For example, the neutral theory explains why rates of amino acid substitutions in proteins are generally consistent with the genomic mutation rates, and why higher substitution rates are observed at sites less involved in protein function [6, 10, 11]. A key observation leading to the development of the neutral theory was the concept of the time-like protein molecular clock. Specifically, the pioneering work of Zuckerkandl and Pauling [2, 3] demonstrated, to the initial surprise of many evolutionary biologists [12], that the number of amino acid substitutions in proteins is approximately proportional to species’ divergence time. It has also been shown that time-like molecular clocks are observed across many different proteins that vary by orders of magnitude in their substitution rates [1, 2, 13].

One of the foundational and still unresolved challenges for the neutral theory and for understanding the origin of protein molecular clock is the generation time effect [14-17]. The generation time effect is the expectation that relatively more protein substitutions should be observed in lineages with shorter generation times than in lineages with longer generation times. As germ-line replication errors are considered the primary source of *de novo* mutations [18, 19], species with longer generation times are expected to accumulate fewer mutations per unit time compared to species with shorter generation times. Due to allele fixation mostly through neutral genetic drift, the rate of protein substitutions should then also be proportional to the number of generations elapsed since species’ divergence from a common ancestor rather than the divergence time itself. But despite a half-century of active research and debate, the role of generation time in protein molecular clock and molecular evolution has not been resolved [4, 20-23]. Although the generation time effect has been observed in plants [24] and invertebrates [25], but among vertebrate species, the generation time effect has been identified for synonymous but not for non-synonymous substitutions [26, 27]. Most importantly, it remains unclear how to quantitatively reconcile the neutral theory prediction of a strong generation effect with the observed molecular clock.

Various mechanisms have been proposed to explain the apparent absence of a strong generation time effect in protein evolution. In particular, Ohta and Kimura proposed an extension of the neutral theory, known as the nearly neutral theory [28, 29]. The nearly neutral theory posits that most accepted substitutions are slightly deleterious rather than neutral, making their fraction and fixation probability negatively correlated with the effective population size. Since species with shorter generation times tend to have larger effective population sizes [30], stronger selection against slightly deleterious alleles may offset, according to the nearly neutral theory, higher mutation rates per unit time in species with smaller generation times and lead to similar substitution rates across species. Although it was indeed observed that smaller fractions of non-synonymous mutations are fixed in species with shorter generation times [31], it is unclear whether this effect can account for the empirically observed patterns of protein molecular substitutions [32]. It was also proposed that an episodic adaptive selection model may explain the absence of the generation time effect [33]. According to this model, proteins undergo rapid bursts of adaptive molecular substitutions followed by long periods of purifying selection [34]. Since these episodic bursts are likely to be driven by adaptation to environmental changes that occur in real time, protein substitution rates should then be independent of the species’ generation time.

In this study, we take advantage of the recently accumulated data on mutation rates across many dozens of mammalian species [35, 36], information about their generation times [37], times of divergence from a common ancestor [38], and available orthologous sequences of protein-coding genes [39] to carefully investigate the relationships between species’ generation time and protein evolutionary rates. We explore how effects such as protein functional and mutation rate selection and the variability of generation times across long evolutionary distances influence patterns of protein evolution. We also develop an integrated model of protein evolution which accounts for these effects. Overall, our work resolves multiple seemingly contradictory observations related to the generation time effect and provides a unifying model which accurately explains the time-like behavior of the protein molecular clock across large evolutionary timescales.

## Results

### Generation time effect in recently diverged sister species

Previous studies, limited by the availability of sequenced genomes, often compared species separated by large phylogenetic distances and with markedly different generation times [11, 27, 40]. However, an important drawback of this approach is that the overall generation time effect gets diluted when analyzing data across long evolutionary branches, as ancestral species likely had different generation times compared to their extant descendants. We addressed this challenge by comparing substitution rates between pairs of sister species (for example, rat-mouse, human-chimp, dolphin-orca) with their corresponding generation times. Because each pair of sister species has only recently diverged, it is very likely that their common ancestors also had similar generation times, which we approximated by the average of the known generation times of extant sister species. In this way, by comparing pairs of sister species with different generation times, and considering their divergence times, we explored the impact of the generation time effect on the rate of molecular evolution.

We selected for our analysis 43 pairs of mammalian sister species that diverged less than 20 million years ago and have similar generation times within each pair (Methods). Using the Orthomam database [39], we identified 2,578 genes whose orthologs are present in all considered species. Using these genes, we then calculated the rate of non-synonymous (*dN*) and synonymous (*dS*) substitutions for each species pair and calibrated these rates with the species divergence times from the TimeTree database [38] (*dN*_*rate*_, *dS*_*rate*_). We observed a strong anticorrelation between *dN*_*rate*_ and species’ generation time (Pearson’s r = −0.69, p <10^−6^, slope = -0.50 in log10-log10 coordinates, Figure 1A) and even stronger anticorrelation between *dS*_*rate*_ and species generation time (Pearson’s r = −0.81, p < 10^−10^, slope = -0.71 in log10-log10 coordinates, Figure 1B). These findings demonstrate, for the first time, a pronounced generation time effect for both synonymous and non-synonymous substitutions across thousands of proteins encompassing all major branches of the mammalian phylogenetic tree.

**Figure 1:**
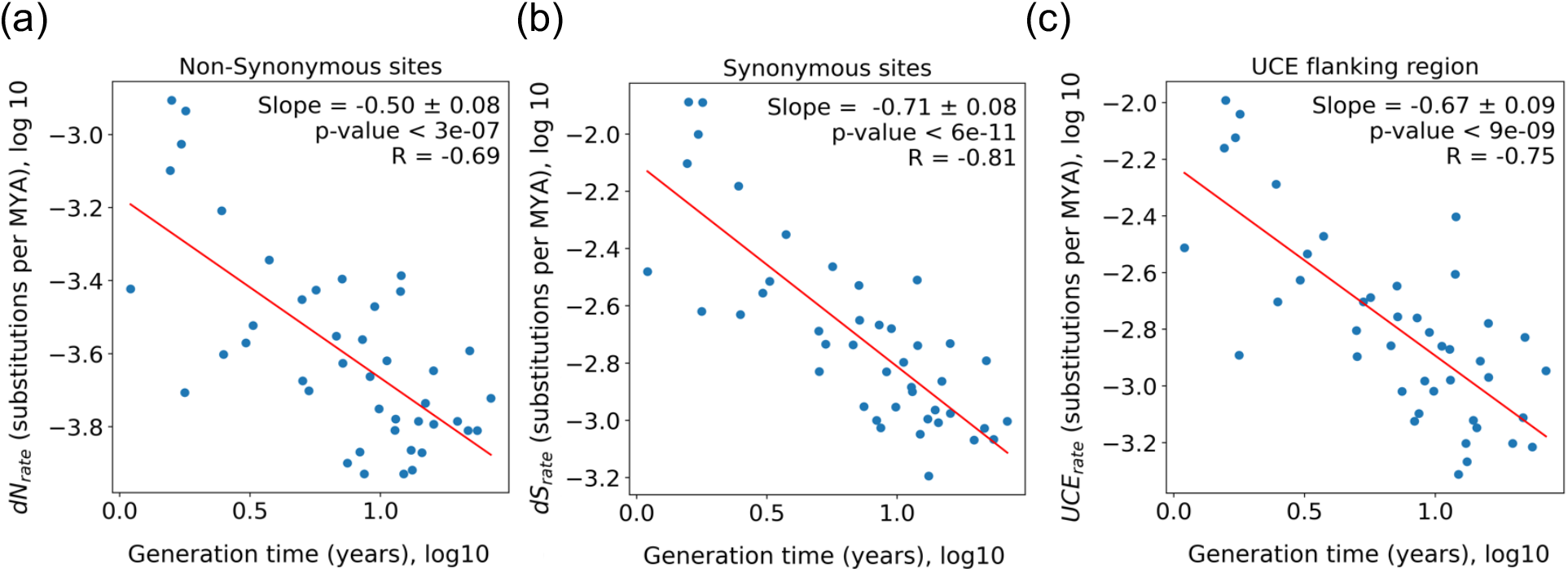
The generation time effect on the rate of evolution in mammalian species. The relationships between per year substitution rates and generation time are shown for (a) non-synonymous sites (*dN*_*rate*_), (b) synonymous sites (*dS*_*rate*_), and (c) UCE flanking regions (*UCE*_*rate*_). Each blue dot (n = 43) represents a pair of mammalian sister species that diverged less than 20 million years ago (MYA). Substitution rates were calculated as the number of substitutions per site accumulated between sister species: (a, b) across 2,578 orthologous proteins or (c) across the flanking sites of 418 UCEs, normalized by the elapsed divergence time (in MYA). The red line represents the linear regression fit. The estimated slopes, the standard error of the slopes, Pearson’s correlation coefficients, and p-values are shown in each figure.

It is widely recognized that much of the non-coding genome evolves neutrally [41, 42], and therefore it should also exhibit a strong generation-time effect. However, unlike protein sites, non-coding regions are rarely highly conserved, making them much more challenging to align. To address this, we focused on ultra-conserved elements (UCEs), which are highly conserved genomic segments (typically 200 to 500 bp in length) found throughout the genome, the majority of which reside in non-coding regions [43, 44]. To investigate the generation time effect in non-coding regions, we used the flanking sequences of these UCEs; these flanking sequences can be aligned due to their proximity to UCEs but they themselves are not under any strong selection pressure. Using baits designed for tetrapods, we identified 418 UCE loci present in all 43 species pairs and extracted the 1000 bp of flanking sequence on both sides of each UCE [45, 46]. We then removed any sites containing more than three gaps and concatenated the outermost 20 bp of these flanking regions from each UCE alignment. This analysis revealed a strong generation time effect in the UCEs flanking regions (Pearson’s r = −0.75, p < 10^−8)^, slope = -0.67 in log10-log10 coordinates, Figure 1C). Notably, the generation time effect scaling observed for the UCEs flanking regions is similar to the one observed at synonymous sites.

We next calculated the correlation between *dN*_*rate*_ and *dS*_*rate*_ for the 43 mammalian sister species and found a very strong relationship (Pearson’s r = 0.96, p < 5 ∗ 10^−24^, Figure 2A). This result suggests that *dN*_*rate*_ substitution rates across sister species align closely with *dS*_*rate*_, but are subject to stronger purifying selection. Similarly, we examined the correlation between *dN*_*rate*_ and the substitution rate at UCE flanking sites (*UCE*_*rate*_) and again observed a high correlation (Pearson’s r = 0.95, p < 8 ∗ 10^−23^, Figure 2B). This finding is striking because UCE flanking sequences lie outside of protein coding regions and therefore should not undergo functional selection related to protein evolution. Their strong correlation with *dN*_*rate*_ suggests that the variability in substitution rates across species is largely driven by life history traits, such as generation time, which affects substitution rates across the entire genome.

**Figure 2:**
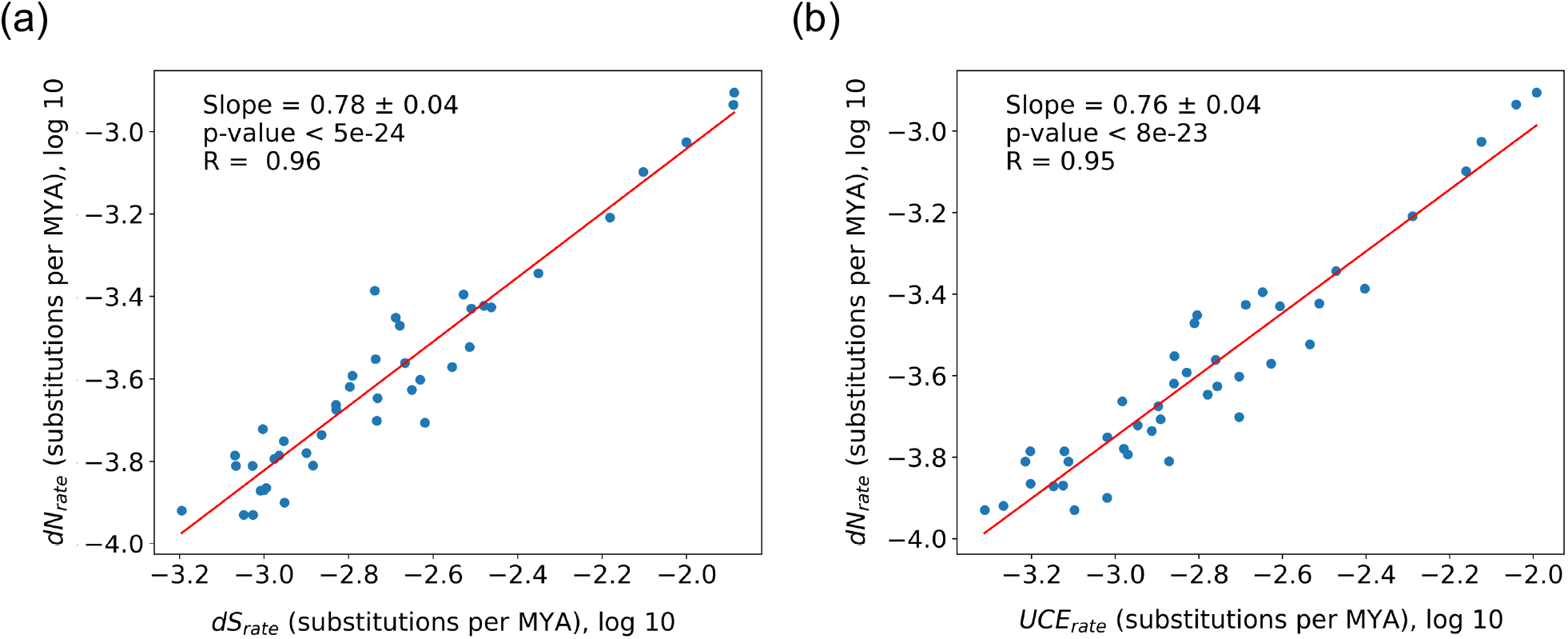
The relationship between substitution rates at different types of sites. The correlation between the substitution rate at non-synonymous sites (*dN*_*rate*_) and the substitution rate at (a) synonymous sites (*dS*_*rate*_) and (b) UCE flanking region sites (*UCE*_*rate*_). Each blue dot represents a pair of mammalian sister species (n = 43). The red line represents the linear regression fit. The estimated slopes, the standard error of the slopes, Pearson’s correlation coefficients, and p-values are shown in each figure.

Notably, empirical data from multiple mammalian species demonstrate a pronounced generation time effect in protein coding regions, both for non-synonymous and synonymous substitutions (*dN*_*rate*_ versus generation time scaling -0.50, and *dS*_*rate*_ versus generation time scaling -0.71). According to the neutral theory of molecular evolution, the fixation of mostly neutral alleles should result in a scaling of -1 between substitution rate and generation time. To reconcile this discrepancy between the predicted and empirically observed scaling, we considered next two effects; the functional selection and the mutation rate selection in species with different generation times.

As Ohta and Kimura proposed [4, 28, 32], slightly deleterious mutations may play an important role in affecting the scaling between *dN*_*rate*_ and generation time due to stronger selection in species with larger effective population sizes. Because generation time and effective population size are negatively correlated [30], species with shorter generation times typically have a lower fraction of mutations that are effectively neutral (*s* < 1*/Ne*), owing to their larger effective population sizes. We investigated the strength of this functional selection effect across mammalian sister species using the ratio of substitution rates at non-synonymous and synonymous sites (*dN/dS*). Consistent with previous studies [31, 32], this analysis demonstrated that the functional selection effect is indeed significant (Pearson’s r = 0.*7*8, p < 10^−9^, Figure 3B) but relatively small in terms of its effect on the scaling (slope of 0.22 in log10-log10 coordinates). Therefore, the selection against slightly deleterious mutations is unlikely to eliminate, as was often suggested [47, 48], the significant generation time effect (Figure 1A).

**Figure 3:**
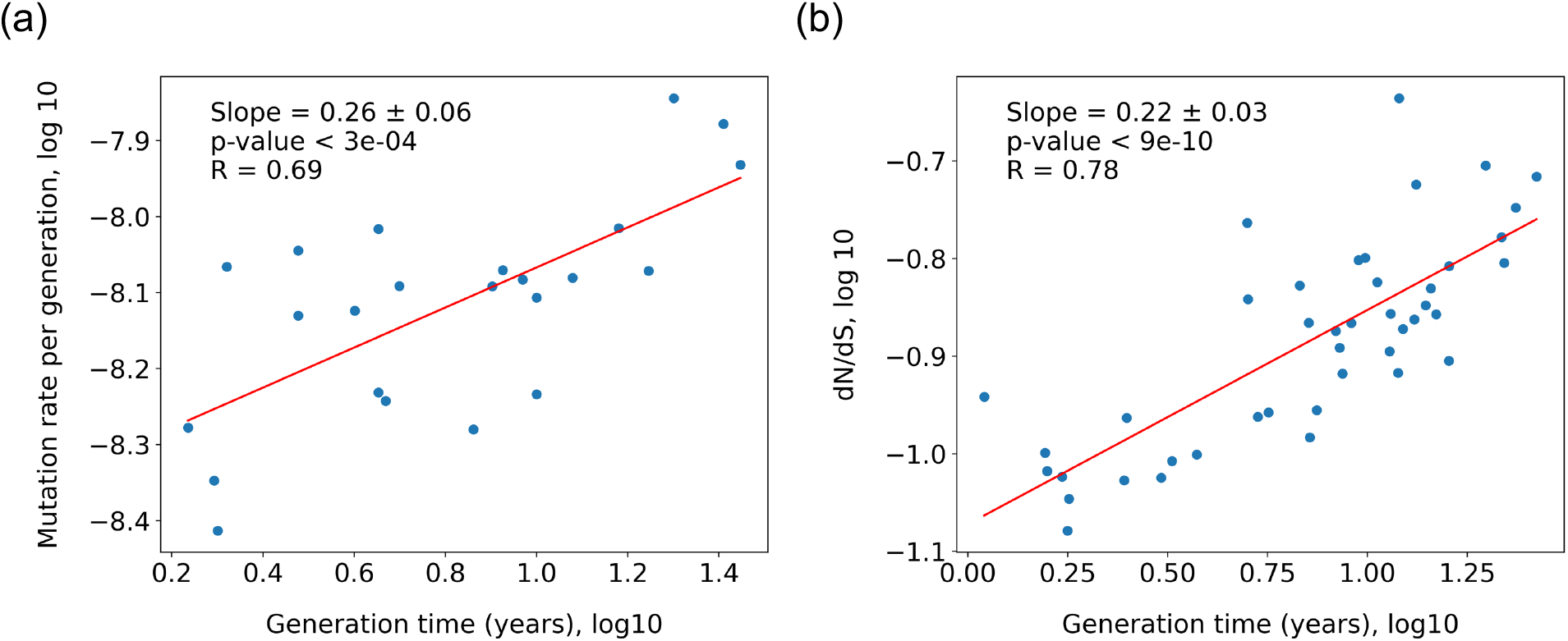
The scaling of mutation rates and the strength of functional selection with generation time in mammalian species. The correlation between generation time and (a) mutation rate per generation (n = 23), and (b) the ratio of non-synonymous to synonymous substitution rates, *dN/dS* (n = 43). Mutation rates were determined from parent-offspring trio sequencing data [35]. *dN/dS* values, which characterize the strength of functional selection (with lower *dN/dS* values indicating stronger purifying selection), were calculated for pairs of mammalian sister species (diverged less than 20 MYA) based on 2,578 orthologous proteins. The red line represents the linear regression fit. The estimated slopes, the standard error of the slopes, Pearson’s correlation coefficients, and p-values are shown in each figure.

Because new mutations are the primary source of fixed substitution, another effect that could influence the scaling between the substitution rates and generation time is the variability of mutation rates across species. Genomic mutation rates have been recently measured across multiple species based on mutation-accumulation experiments [36] and through parental-offspring trio sequencing [35, 36]. These experiments demonstrate that species with smaller population sizes [36] or longer generation times [35] generally exhibit higher mutation rates. These observations support the hypothesis that species with larger effective population sizes experience stronger selection pressure to reduce their mutation rates, thereby decreasing the overall mutational load [49, 50]. Additionally, higher mutation rates per generation in long-lived species may arise from a greater number of germline cell divisions within a single generation [51]. To examine how mutation rates depend on generation time, we analyzed mutations rates across 23 mammalian species from a recent parent-offspring sequencing study [35]. Our results reveal a significant correlation between mutation rate and generation time (Pearson’s r = 0.69, p < 10^−3,^, Figure 3A), with a log10-log10 slope of 0.26.

Considering both functional and mutations rate selection makes the predicted scaling of the generation time effect in evolution of proteins and UCE flanking sequences consistent with empirical observations. Specifically, a reduction of mutation rate in species with smaller generation times is expected to decrease the scaling between the rate of substitutions for mostly neutral mutations and generation time to -0.74 ± 0.06 (mean ± standard deviation), in line with the empirically observed generation time scaling for synonymous protein sites -0.71 ± 0.08 and for UCE flanking region sites -0.67 ± 0.09. Similarly, for non-synonymous sites, the empirically observed generation time scaling of -0.50 ± 0.08 matches the theoretically expected scaling of - 0.52 ± 0.09 once both the mutation rate selection and functional selection effects are considered. Overall, these results demonstrate that the observed variability in substitution rates with generation time for non-synonymous, synonymous and UCE flanking sites in mammalian species is consistent with the nearly neutral theory of molecular evolution when key genomics and functional effects are included.

### The influence of protein conservation on the generation time effect

Protein evolutionary rates vary substantially among different proteins in the same species [13, 52]. Given that mutation rate remains relatively constant across the genome [53], the variability of functional selection between proteins may influence the strength of the generation time effect. To investigate this, we ranked proteins based on their evolutionary conservation score using *dN* between human and mouse (see Methods) and calculated the scaling of *dN*_*rate*_, *dS*_*rate*_, and *dN/dS* with generation time as a function of protein conservation. Interestingly, we found that for non-synonymous sites, the generation time effect becomes stronger for less conserved proteins (see Figure 4A). Moreover, this observation confirms that the generation time effect is affected by the fraction of sites with weak functional effects that are more prevalent in less conserved proteins. We also found that the functional selection effect, quantified by the scaling of *dN/dS* with generation time, is more pronounced for conservative proteins (see Figure 4C). Reassuringly, the generation time scaling for the synonymous sites, *dS*_*rate*_, substitutions remain constant across all conservation levels (Figure 4B), consistent with the expectation that synonymous sites are not strongly affected by functional selection.

**Figure 4:**
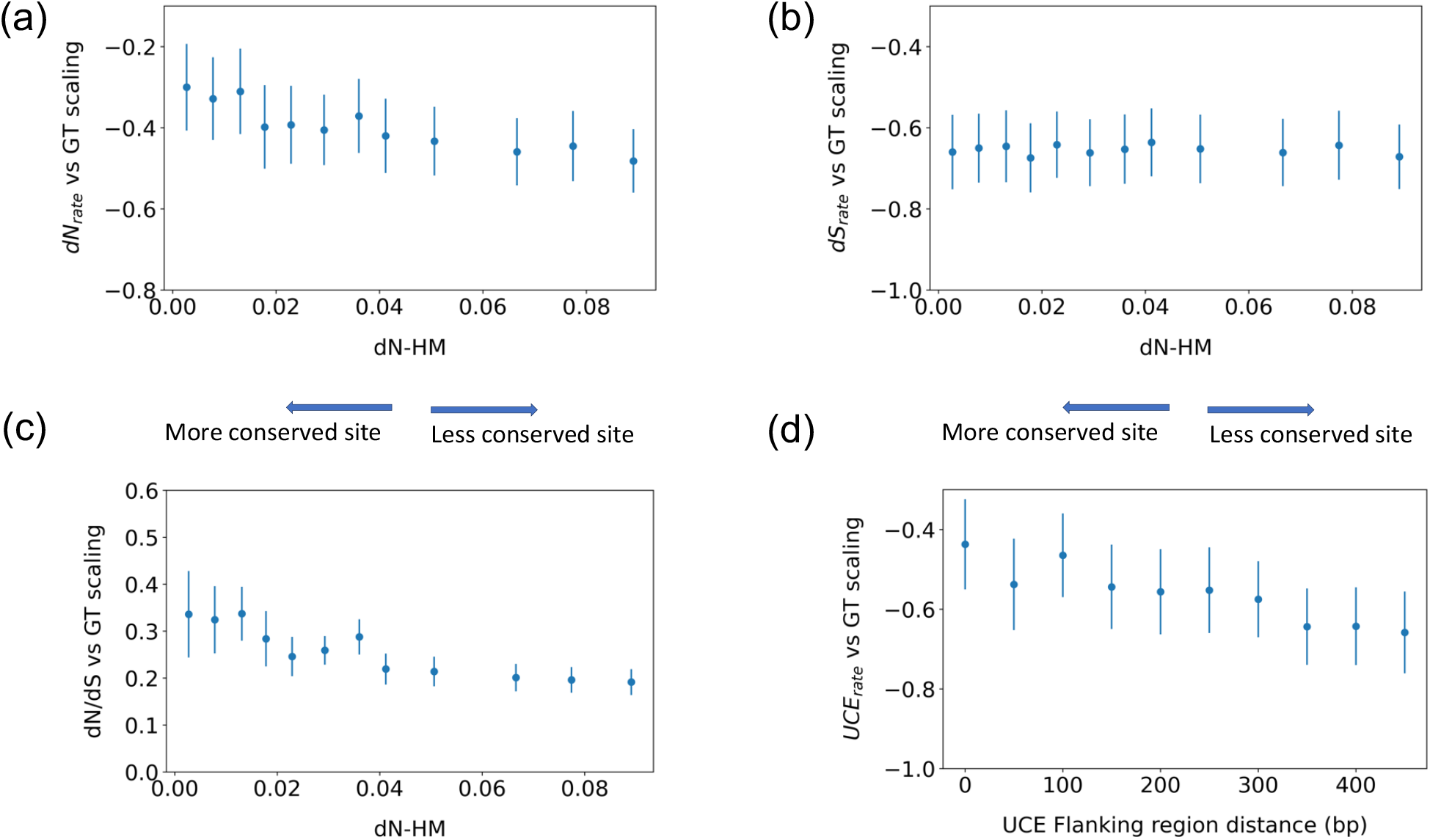
The influence of proteins and UCE flanking sites conservation on the strength of the generation time effects. The slopes of the relationships between generation time and (a) substitution rates at non-synonymous sites, (b) substitution rates at synonymous sites, (c) functional selection measured by the *dN/dS* ratio, and (d) substitution rates at UCE flanking regions are shown in relation to the level of sequence conservation. The X-axes represent (a, b, c) the number of non-synonymous substitutions (per site) accumulated between human and mouse, and (d) the distance from the UCE flanking region segment to the UCE boundary (defined as 500 bp distance from the UCE). Each point represents a bin of 200 proteins with similar conservation levels (a, b, c), or UCE flanking region sites of 20 bp in length (d) The protein or UCE sequences within each bin were concatenated and the scaling slopes corresponding to each bin calculated. Error bars represent the standard error of the slopes corresponding to each bin.

Similarly to protein-coding regions, the generation time effect is likely to vary within the UCEs flanking regions, as flanking sequences closer to UCE boundaries should experience relatively stronger purifying selection compared to more distant sections of the flanking regions. To investigate this effect, we calculated the generation time effect scaling for different sub-regions of the UCE flanking sequences as a function of their distance to the UCE boundary (Figure 4D). To avoid including the UCEs themselves, we analyzed only those flanking regions located at least 500 bp away from the edge of any UCE. Interestingly, we found that the generation time scaling increases almost linearly as one moves farther from the UCE boundary. Flanking sequences most distant from UCEs generally exhibited a stronger generation-time effect, approaching the one observed for synonymous sites. In contrast, flanking sequences closer to UCEs exhibit weaker generation times scaling, more similar to the scaling of non-synonymous sites.

### Consistency of the generation time effect with the time-like molecular clock

Our analysis of recently diverged (< 20 million years) sister species revealed a pronounced generation time effect for both non-coding and protein coding sites (Figure 1). Moreover, when several effects, such as functional and mutation rate selection, are considered, the predicted generation time effect agrees well with empirical observations and expectations form the neutral theory. However, over longer evolutionary periods, species descending from a common ancestor are unlikely to maintain similar generation times. Thus, to investigate the generation time effect and develop a comprehensive model of protein evolution at longer timescales, we compared accumulated substitutions across 200 proteins among all 903 unique possible pairs of the 43 mammalian species considered in our analysis. To that end, we used a ratio-based approach that does not depend on species’ divergence time estimates, but instead computes the ratio of amino acid substitutions accumulated by two species since their divergence from a common ancestor to the ratio of their present generation times (see Methods) [40]. Due to the generation time effect, species with shorter generation times should accumulate relatively more substitutions during their divergence than those with longer generation times. Consistent with this prediction, we observed a significant correlation between the ratio of species’ generation times and the ratio of accumulated substitutions (Pearson’s r = -0.72, p < 10^−15^, Figure 5C), with a slope in log10-log10 coordinates of -0.28 ± 0.01. This result demonstrates that the generation time effect remains significant even at longer evolutionary timescales.

**Figure 5:**
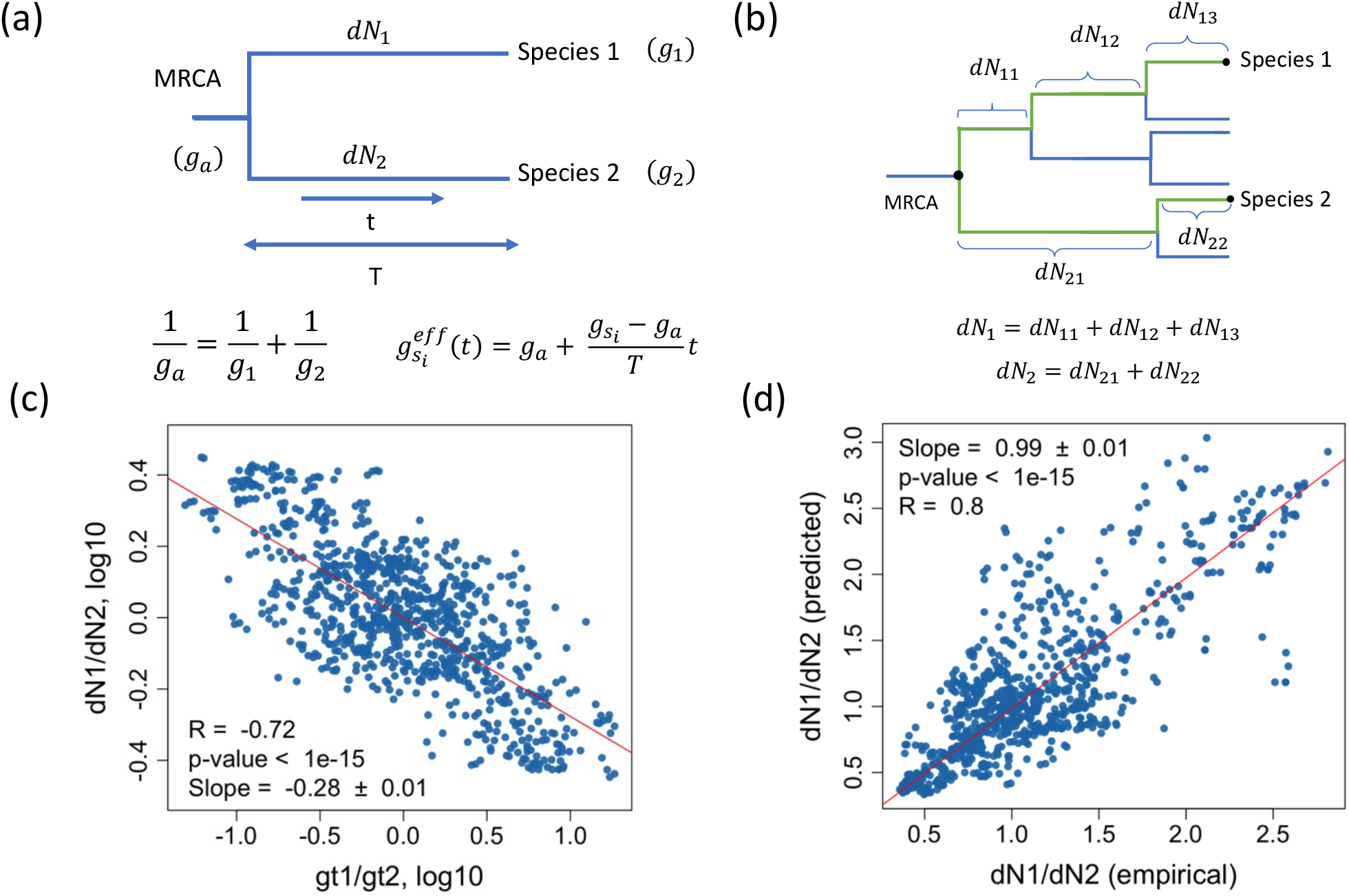
The integrated model of the protein molecular clock across long evolutionary timescale. (a) The generation time of the common ancestor (g_*a*_) was modeled as the harmonic mean of the daughter species’ generation times. The effective generation time of a species 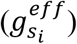 was assumed to change linearly with time from the ancestor’s generation time to its current value. (b) The model was sequentially applied to estimate the generation times of ancestral nodes and to calculate the number of substitutions for intermediate branches. (c) The correlation between the empirically observed ratio of the numbers of substitutions at non-synonymous sites accumulated during independent evolution of two daughter species and the ratio of their generation times. (d) The correlation between the empirically observed substitutions ratio and the ratio predicted by the integrated model of the molecular clock (Methods). Each blue dot represents a unique pair of species (n = 903). The red lines represent the linear regression fit. The estimated slopes, the standard error of the slopes, Pearson’s correlation coefficients, and p-values are shown.

Next, to account for the changes of generation time along the evolutionary branches (since divergence from a common ancestor), we employed a simple model where the generation time of a species changes linearly from the most recent common ancestor to its current value (Figure 5A, see Methods). The generation time of the common ancestor was calculated by taking the harmonic mean of the generation times of its daughter species. To estimate the generation times of all previous ancestors of the considered species, we applied this approach sequentially to all ancestral nodes in the phylogenetic tree (Figure 5B), (see Methods). The advantage of this model is that the empirically established scaling of functional and mutation selection with generation time (Figure 3) makes it possible to incorporate these two important effects into the model. By using the integrated model, which includes the combined effects of the generation time changes along a phylogenetic branch, functional selection, and mutation rate selection, it is then possible to calculate the expected numbers of protein substitutions along any evolutionary branch (see Methods) (Figure 5A).

Using the integrated model, we calculated the expected ratio of accumulated non-synonymous protein substitutions (*dN*_1_*/dN*_2_) between all pairs of species, including species with very different generation times. Despite its simplicity, we found that the integrated model predicts remarkably well the empirically observed differences in the ratio of accumulated substitutions (Pearson’s r = − 0.8, *p* < 10^−15.^, slope = 0.99 ± 0.01, Figure 5D). We note that the slope between the predicted and empirically known *dN*_1_*/dN*_2_ ratios is very close to one, indicating that the model accurately captures the magnitude of generation time effect along deep evolutionary branches.

The protein molecular clock is often supported by the observation that divergence between orthologous sequences increases in an approximately linear fashion with the time since species’ divergence [1, 6, 11]. We next investigated whether our integrated molecular clock model is consistent with that observation by examining the divergence of 323 protein sequences between human and 177 other mammals those orthologous sequences are available in the Orthoman database [39]. In agreement with previous results [47], we observed that the average sequence divergence for non-synonymous protein sites largely exhibits a linear divergence with time (Figure 6A, blue dots), even though the considered species differ by more than an order of magnitude in their generation times. We then used our integrated clock model, which accounts for mutation rate selection, functional selection, and generation time variability, to predict sequence divergence over time for the same set of species (Figure 6A, red dots). We found an excellent agreement between the model’s prediction and the empirically observed sequence divergence across 160 million years of mammalian evolution (Pearson’s r = 0.95, p < 10^−15.^). The model not only predicted the overall linear trend in sequence divergence but also captured the residual deviation from that trend (Pearson’s r = 0.81, p < 10^−15.^, slope = 0.93 ± 0.05, Figure 6B). These deviations arise mostly from generation time effects; for example, the sequence divergence of primates (with long generation times) is below the overall linear trend, whereas the sequence divergence of rodents (with short generation times) is above the trend. Notably, the residual slope (Figure 6B) is nearly equal to unity, confirming again that the model can accurately capture the small deviations of sequence divergence driven by differences in generation time

**Figure 6:**
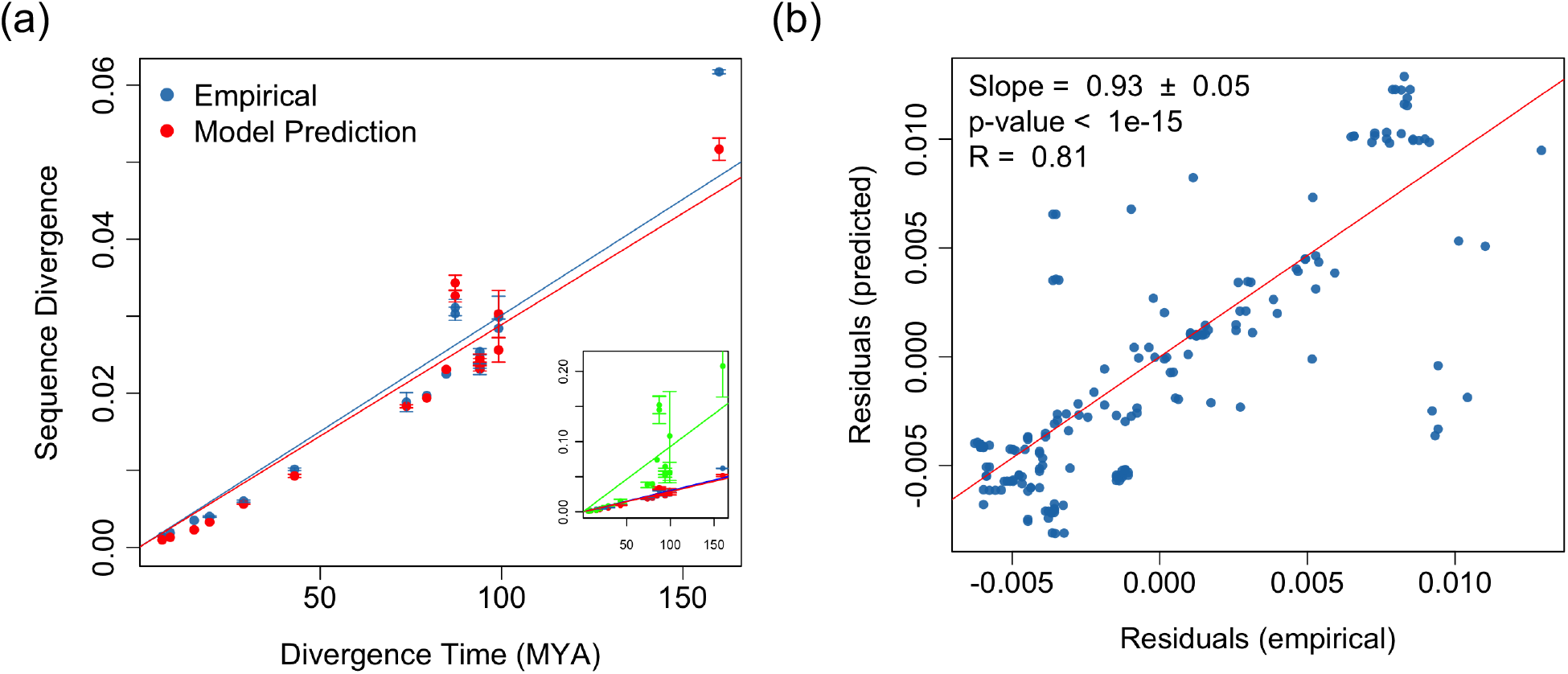
Integrated model of the molecular clock applied to sequence divergence between human and mammalian species. (a) Sequence divergence at non-synonymous sites, *dN*, between human and multiple mammalian species (n = 177) as a function of divergence time. Blue dots represent empirical data, and red dots represent the model prediction. Each dot represents the average sequence divergence across 323 proteins and all species that diverged from human at the corresponding time point. The error bars indicate the standard deviation. In the insert, green dots represent sequence divergence at synonymous sites. The blue, red and green lines show the linear regression fits to the corresponding data (b) The correlation between the empirical residuals from the overall linear trend and the residuals predicted by the integrated clock model. Residuals were calculated as the deviation from the linear regression fit of empirical sequence divergence (blue line in panel (a)). Each blue dot represents a mammalian species (n = 177). The red line represents the linear regression fit. The estimated slopes, the standard error of the slopes, Pearson’s correlation coefficients, and p-values are shown.

## Discussion

The results presented in our paper reconcile multiple, seemingly contradictory observations related to protein evolution across species. According to the neutral theory [5, 7], most substitutions in proteins are effectively neutral with respect to a species’ fitness. But if these substitutions are neutral, one would expect a pronounced generation time effect, such that species with shorter generation times accumulate relatively more substitutions per unit time than species with longer generation times. While a strong generation time effect is frequently observed for synonymous substitutions across diverse taxa, detecting this effect for non-synonymous substitutions in vertebrates has been challenging [26, 27], thus calling into question key tenants of the neutral theory. In response, researchers have proposed, over the past 50 years, various mechanisms to explain the apparent absence of a strong generation time effect for amino acid substitutions. In particular, the nearly neutral theory [29] suggested that stronger selection against slightly deleterious protein-coding mutations in species with larger population sizes and shorter generation times may explain the absence of a pronounced generation time effect.

Notably, in one of his latest papers Kimura emphasized the need for additional experimental data to resolve the issue of the generation time effect and its relationship to the neutral theory [54]. Following this suggestion, in this study we took advantage of the recently accumulated evolutionary and genomic data in mammalian species [35, 36, 39]. Our analyses reveal, for the first time in vertebrates, a pronounced generation time effect. We also found that the mechanism proposed by the nearly neutral theory, i.e., stronger functional selection in species with shorter generation times, has a significant impact but plays a relatively minor role in reducing the generation time effect. Furthermore, when considering both functional selection and mutation rate selection, the generation time effect we observe is fully consistent with predominantly neutral protein substitutions. We note that our results do not dismiss the influence of various adaptive phenomena, such as hitchhiking and background selection in proteins evolution [55]. Nevertheless, they suggest that over long evolutionary timescales protein evolution is consistent with the fixation of mostly neutral substitutions, with the substitution rate primarily driven by the mutation rate.

Another longstanding puzzle in molecular evolution is how to explain the apparent time-like behavior of the protein molecular clock observed across species with vastly different generation times [11]. To address this question, we developed an integrative model that traces changes in a species’ generation time along its evolutionary history and calculates how these shifts influence the accumulation of substitutions. Our findings resolve the clock paradox, demonstrating that the protein clock is fully consistent with an integrated framework. Notably, we show that one of the major contributors to the time-like property of the protein clock is the gradual and continuous changes in generation time as species diverge from a common ancestor. While there is a substantial short-term variability in the clock rate, as seen in our analysis of closely related species, this variability diminishes over longer evolutionary timescales, resulting in the emergence of time-like behavior.

Although the results presented here capture the behavior of protein molecular clock across approximately 160 million years of mammalian evolution, it would be interesting in the future to extend similar analysis to include both vertebrate and invertebrate species. In particular, it will be insightful to investigate how the various evolutionary and genomic effects considered in our analysis affect the clock rate on longer timescales. Moreover, the model we developed can be extended by integrating data on species’ life histories derived from fossil records [56-58]. In recent work, we have shown how the variability of evolutionary rates between different proteins in the same species may emerge from differences in functional optimization, such as the variability in enzymatic kinetic constants, necessary to minimize expression costs in specific tissues and cell types [59, 60]. We found that stronger functional optimization intensifies purifying selection in highly expressed proteins, thus slowing their evolutionary rates. Combining the life history effects examined in this study with structural constrains and functional optimization could pave the way for an integrated functional theory of protein evolution, capable of describing protein evolution both within and between species.

## Methods

### Species generation time

The generation time for species in the analysis were obtained from a database containing information on extant mammalian species [37]. For the sister species analysis, the generation time for each pair of sister species was calculated by averaging the generation times of both species. When species’ generation time was unavailable in the database, we used the average generation time of the corresponding clade. If data for the entire clade were not available in the database, we used the generation time of the other sister species in the pair.

### Calculation of evolutionary rates for sister species

We constructed an orthologous gene dataset using data from the OrthoMam database [39], which provides alignments of coding sequences (CDS) for one-to-one orthologs across 190 mammalian species. For our analysis, we selected phylogenetically independent pairs of sister species based on the following criteria: (i) the sister species diverged at least 2 million years ago, ensuring that a significant number of non-synonymous substitutions have accumulated between them, and (ii) the divergence time is less than 20 million years, ensuring similar generation times in both species. Using these criteria, we identified 43 pairs of sister species. Next, we selected genes whose orthologs were present in each of the considered species, resulting in a dataset of 2578 orthologous mammalian genes.

To analyze the overall generation time scaling, we concatenated the OrthoMam alignments for selected genes into a single alignment containing over 5 million nucleotides. For each sister species pair, we then calculated the rates of synonymous (*dS*) and non-synonymous (*dN*) substitutions, as well as the *dN/dS* ratio using Codeml (runmode =-2) from the PAML package [61]. To obtain per year substitution rates (*dS*_*rate*_ and *dN*_*rate*_), we divided the corresponding values, *dS* and *dN*,by the divergence times between sister species, based on the TimeTree database [38].

For analysis of the influence of protein conservation on the generation time effect, we first ranked proteins based on their conservation. For this we used the rate of non-synonymous substitutions between human and mouse orthologs, *dN*^*Human-Mouse*^, calculated with Codeml [61] as a representative evolutionary rate. We then grouped proteins, ranked from the most conserved to the least conserved, into bins of 200 proteins each and concatenated the corresponding alignments within each bin into a single alignment. Next, for each bin, we calculated *dS, dN*, and *dN/dS* using Codeml [61] and calculated their scaling with generation time.

### Analysis of UCE flanking sequences

We downloaded the reference genomes for all 43 pairs of sister species from the NCBI website. To extract the UCE loci from these genomes, we used 5,472 baits targeting 5,060 UCEs in tetrapods [45]. Using the Phyluce software [62], we located and extracted the UCE loci from the genomes along with 1000 bp flanking regions on both sides. We kept only the UCE loci that were present in every species under consideration and aligned them using MAFFT [63], resulting in a total of 418 UCE alignments. To calculate the number of substitutions in UCE flanking sites between the sister species, we selected 20 bp from the outer portion of the flanking regions of each UCE alignment after removing the sites more than 3 gaps and concatenated them into a single alignment. For this alignment we then inferred phylogenetic tree using IQ-TREE [64] and summed up the lengths of branches connecting sister species through their common ancestor.

We also investigated how the generation time scaling is influenced by the distance between the segment of UCE flanking region and the UCE boundary. To avoid including the UCEs themselves, we analyzed only the portions of the flanking regions located at least 500 bp away from the edge of any UCE. Starting from this position, we selected the first 20 bp of each UCE flanking sequence after removing the sites with 3 gaps, calculated substitution rates between sister species as described above, and obtained the generation time scaling for these segments. Then, we shifted the focus segments outward by 50 bp and repeated the procedure until reaching the outer boundaries of UCE flanking regions.

### Deep evolutionary tree analysis

For the analysis of the generation time scaling across deep evolutionary branches, we selected one species from each of 43 pairs of sister species. Additionally, we included *Ornithorhynchus anatinus*, which diverged from *Homo sapiens* approximately 170 million years ago, as a common outgroup to all other mammalian species. The phylogenetic tree of the resulting 44 species was constructed using IQ-TREE software [64], which utilizes maximum likelihood estimations to determine phylogenetic relationship between the species. The constructed phylogenetic tree was then used in Codeml (model = 1, runmode = 2) to calculate the number of non-synonymous substitutions per non-synonymous site (*dN*) for each branch of the phylogenetic tree [61]. Then, for every possible pair of species (excluding *Ornithorhynchus anatinus*), we calculated the total number of substitutions (per site) for both species from their common ancestor by summing the *dN* values along the intermediate branches leading to the common ancestor. Finally, for each pair, we calculated the substitution ratio (*dN*_1_*/dN*_2_) and plotted it against their generation time ratio (g*t*_1_*/*g*t*_2_).

### Sequence divergence between human and other mammals

For the analysis of long evolutionary sequence divergence, we used all mammalian species available in the OrthoMam database [39], for which generation time data were available [37], resulting in a total of 177 species. We selected genes with orthologs present in all 177 species, yielding 323 orthologous genes. The OrthoMam alignments for the selected genes were concatenated into a single alignment, and the non-synonymous sequence divergence between human and other species was calculated using the pairwise mode of Codeml [61]. Corresponding divergence times were obtained from the TimeTree database [38].

### Integrated model of protein molecular clock

According to the neutral theory [6], at neutral sites, the substitution rate equals the mutation rate (μ). Furthermore, at other sites, the substitution rate is scaled down by a factor corresponding to the proportion of neutral mutations at these sites. For non-synonymous sites in protein-coding genes, this factor can be estimated using the ratio of non-synonymous to synonymous substitution rates (*dN/dS*) [65]. Thus, the rate of non-synonymous evolution can be expressed as:

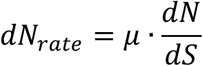

Because we are interested in the influence of species’ generation time on the evolution of their proteins, we further express the mutation rate and *dN/dS* ratio as functions of generation time, using the scaling obtained from the sister species analysis (Figure 3). Specifically, the mutation rate per generation was expressed as 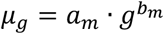, where *lo*g_10_(*a*_*m*_) and *b*_*m*_ are the intercept and slope obtained from fitting the dependence of the mutation rate (per generation) on generation time in log_10_-log_10_ coordinates (Figure 3A). Similarly, 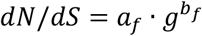, where *lo*g_10_(*a*_*f*_) and *b*_*f*_ are the intercept and slope obtained from the fitting the dependence of *dN/dS* on generation time in log_10_-log_10_ coordinates (Figure 3B). Therefore, the substitution rate can be expressed as a function of generation time as follows:

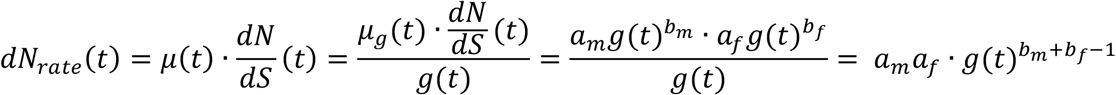

We assumed that the generation time of a species changes linearly between the most recent ancestor and the current value: g(*t*) = g_*a*_ + (g_*s*_ − g_*a*_) · *t/T* where g_*a*_ is the generation time of the ancestor node, g_*s*_ is the generation time of the species, T is the time elapsed since their divergence, and *t* is the time passed since the ancestor node. The integration over the elapsed time gives the number of accumulated substitutions in the branch in terms of generation times of the species and the ancestor as follows:

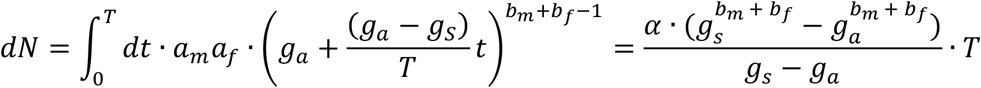

where 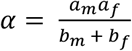. The values of the coefficients were obtained from the fit of empirical data, and are as follows *a*_*m*_ = 10^−8.31)^, *a*_*f*_ = 10^−1.07^, *b*_*m*_ = −0.26 and *b*_*f*_ = −0.22.

We further applied the integrated model of the molecular clock to describe sequence divergence across the deep evolutionary branches. To this end, we first obtained the divergence times for each branch of the phylogenetic tree of the 177 mammalian species used in our analysis, utilizing the TimeTree database [38]. Then, we estimated the generation time of ancestral nodes using the weighted harmonic mean of the generation times of the daughter nodes, where the weight was the inverse of the time elapsed between each daughter node and their common ancestor. Once both the generation time for each node and the divergence times between them were calculated, we used our integrated model to compute the number of substitutions (per site) along each branch. For each pair of species used in the analysis of the generation time effect (Figure 5C), we estimated the expected ratio of the numbers of substitutions accumulated in each species from their last common ancestor (Figure 5D). To this end, for each species, we summed the predicted number of substitutions of all intermediate branches between the species and the common ancestor. Notably, as we were interested in the ratio of the number of substitutions, the predicted value of the ratio depended only on the constants *b*_*m*_ and *b*_*f*_, which characterize the slopes of the scaling of the mutation rate and *dN/dS* with the generation time.

To predict sequence divergence between human and other mammalian species (Figure 6), for each species, we summed the predicted number of substitutions of all intermediate branches of the tree between human and the common ancestor node and between species and that ancestor. In this analysis, we considered the absolute number of substitution (per site); thus, the model predictions depended on all parameters of the model, including both the slopes and the intercepts of the scaling of the mutation rate and *dN/dS* with the generation time.

## Acknowledgments

This work was supported in part by the National Institute of General Medical Sciences Grant No. R35GM131884 to D.V.

